# Cryo-EM structures of α-Synuclein(31-100) amyloid fibrils reveal disease-like structural motifs without reproducing the Parkinson’s Disease polymorph

**DOI:** 10.64898/2026.07.20.739490

**Authors:** Kristina Biedermann, David Rhyner, Lukas Frey, Roland Riek, Jason Greenwald

## Abstract

The structural diversity of alpha-synuclein amyloid fibrils is closely linked to the pathogenesis of Parkinson’s disease and related synucleinopathies. However, reproducing disease-associated fibril conformations from recombinant full-length protein *in vitro* has remained challenging. Inspired by successful truncation strategies developed for the Tau protein, we investigated whether removing the disordered terminal regions («fuzzy coat») of alpha-synuclein could bias fibril assembly toward disease-relevant folds. We designed a truncated construct comprising residues 31-100, corresponding to the structured core of patient-derived Parkinson’s disease fibrils, and systematically screened aggregation conditions across a broad range of pH values and ionic environments. Cryo-electron microscopy revealed four previously undescribed fibril structures, including new subtypes of the established type 1 and type 3 polymorphs and a novel fibril fold, termed type 10, which reproducibly formed under acidic conditions. Type 10 was observed as two distinct dimeric assemblies (10A and 10B) that share a common protofilament fold but differ in their inter-filament interfaces. Structural comparison with the patient-derived Parkinson’s disease polymorph revealed local similarities, including conserved β-strand organization and loop conformations within the fibril core, but remains structurally distinct overall. Our results demonstrate that rational construct design combined with systematic environmental screening reshapes the alpha-synuclein polymorphic landscape and promotes structural motifs characteristic of disease-associated fibrils.

## Introduction

Alpha-synuclein (αSyn) is an intrinsically disordered protein that remains largely monomeric under physiological conditions.^1,2^ αSyn can aggregate into highly ordered amyloid fibrils which are characterized by the cross-β sheet motif.^3,4^ The aggregation process is believed to proceed via the amyloid-typical nucleation-dependent polymerization pathway, including a secondary nucleation mechanism, where existing fibrils catalyze the formation of new nuclei and thereby accelerate fibril proliferation.^5^ This aggregation process with the formation of amyloids is a central pathological hallmark of multiple neurodegenerative disorders, such as Parkinson’s disease (PD) and related synucleinopathies.^6–8^ It has become apparent that αSyn fibrils are not structurally uniform. Instead, they exist as distinct polymorphs arising from multiple local minima in the aggregation energy landscape and from kinetic trapping during fibril assembly.^9^ Importantly, the structural diversity of αSyn fibrils extends beyond biophysical variation and correlates with distinct disease phenotypes, supporting the notion that individual polymorphs encode pathogenic information at the molecular level.^9–12^

Advances of cryo-electron microscopy (cryo-EM) have fundamentally transformed our understanding of the structural landscape architecture of amyloids at atomic resolution. High-resolution structures of *ex vivo* derived αSyn fibrils have revealed that disease-associated polymorphs possess unique protofilament folds and intermolecular interfaces.

However, a persistent challenge remains: disease-associated αSyn polymorphs identified in *ex vivo* material cannot currently be reliably reproduced by *in vitro* aggregation of recombinant αSyn.^13^ Full-length recombinant αSyn(1-140) readily forms amyloid fibrils under a wide range of *in vitro* conditions, with environmental factors such as pH, sample purity and agitation protocol strongly influencing polymorph formation.^11,12,14–18^ Nevertheless, the resulting fibrils generally differ structurally from those isolated from patient tissue. Thus, a substantial gap remains between *in vitro* aggregation and *in vivo* pathology.^10,15^

Posttranslational modifications and proteolytic processing have emerged as important modulators of αSyn aggregation. In particular, N- and C-terminal truncations are frequently observed in pathological inclusions and have been shown to modulate aggregation kinetics and fibril structure.^19–21^ A compelling precedent exists in the Tau field, where removal of flexible terminal regions enabled the successful *in vitro* reconstruction of disease-relevant folds characteristic of Alzheimer’s disease and chronic traumatic encephalopathy. Notably, Lövestam et al. demonstrated that rational truncation strategies, combined with systematic screening of solution conditions (i.e. ionic strength, buffer, additives), can drive the formation of specific pathological conformations.^22^ Inspired by this approach, we aimed to explore whether an analogous strategy could be applied to αSyn. Guided by structural insights from *ex vivo* PD fibrils, we designed a truncated construct that lacks the «fuzzy coat». Accordingly, it includes only the residues in the ordered core, as identified in high-resolution cryo-EM structures.^11^ This results in a construct spanning residues 31-100 (αSyn(31-100)), thereby excluding both the N-terminal amphipathic region and the acidic, negatively charged C-terminal tail, which are largely unresolved in patient-derived structures. The design aims to reduce the accessible conformational landscape and bias the aggregation pathway towards the structurally defined core of the PD polymorph.

Within this framework, we determined the molecular structure of a novel fibril polymorph, termed type 10. This polymorph exhibits partial structural similarity to the fold associated with PD and forms reproducibly under acidic conditions (pH 4). Although it shares characteristic secondary-structure elements and local conformational motifs with the disease-associated polymorph, the type 10 polymorphs overall still differ from the PD polymorph. Moreover, we identified two polymorph subtypes, 1t and 3F, that were previously observed only for full-length αSyn. Together, these findings indicate that targeted manipulation at the protein sequence and physicochemical conditions can drive αSyn aggregation while a recapitulation of the PD specific polymorph remained elusive.

## Results and Discussion

### Environmental conditions shape the polymorphic landscape of αSyn(31-100)

To investigate how environmental conditions influence fibril formation of the αSyn(31-100) construct, aggregation conditions were screened across a broad range of pH values and ionic conditions (Table 1). From this screen, four previously unreported fibril polymorphs were identified (Fig. 1). Two of these structures correspond to new subtypes of already described type 1 and type 3 polymorphs termed type 1t and 3F, whereas two represent a distinct fibril fold observed in two subtypes, termed type 10A and type 10B following our naming scheme.^15^

**Figure 1.**
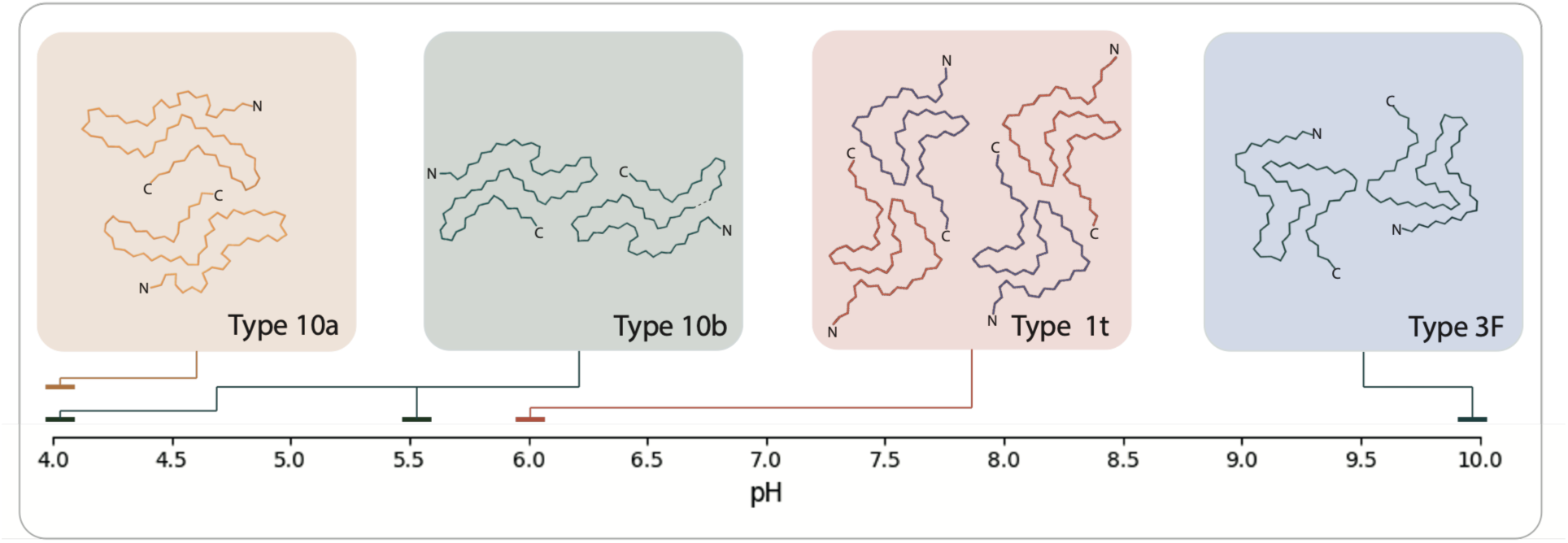
The polymorphic landscape of αSyn(31-100). A) Overview of fibril polymorphs identified across the screened aggregation conditions. The horizontal axis indicates the pH range under which the corresponding polymorphs were observed. The backbone trace of a single layer within the fibril is shown for each of the polymorphic structures.

**Table 1:** Aggregation conditions for αSyn(31-100): Overview of aggregation conditions screened for the truncated αSyn(31-100) construct.

| Protein concentration [μM] | pH | Buffer | Ionic strength [mM], Salt, Additives | Polymorph |
| --- | --- | --- | --- | --- |
| 40 | 10 | Borate-phosphate | - | 3F |
| 20 | 6 | PBS | 200 mM MgCl<sub>2</sub> | 1t (tetramer) |
| 20 | 4 | 0.1 M Citrate | 200 mM CaCl<sub>2</sub> | 10A |
| 20 | 4 | 0.1 M Citrate-phosphate | 200 mM MgCl<sub>2</sub> | 10A |
| 20 | 4 | 0.1 M Citrate-phosphate | 200 mM MgCl<sub>2</sub><br>5 mM Poly(A)<br>(average chain length: 60) | 10A |
| 20 | 5.5 | 0.1 M Citrate-phosphate | 200 mM MgCl<sub>2</sub> | 10B |
| 20 | 4 | 0.1 M Citrate | 200 mM NaCl | 10B |
| 20 | 4 | 0.1 M Citrate-phosphate | 200 mM NaCl | 10B |
| 20 | 4 | 0.1 M Citrate-phosphate | 200 mM KCl | 10B |

The identified structures exhibit a strong dependence on aggregation conditions. At pH 6 in the presence of 200 mM MgCl_2_, a tetrameric subtype of the type 1 polymorph was observed (Fig. 2A-D). Representative cryo-EM data and validation are shown in the Supplementary (Fig. S3). In contrast to the already reported type 1 architectures, the fibril assembles into a four-protofilament arrangement with two distinct two-fold symmetric protofilament interfaces. Inspection of the smaller protofilament interface (Fig. 2C) suggests a stabilization through electrostatic interactions between Lys58 and Glu83 providing two symmetric charged interactions typical of those observed in other αSyn protofilament interfaces. In striking contrast, the larger and extended (and again two-fold symmetric) protofilament interface covers both hydrophobic interactions between Phe94 and Val66, a salt bridge between Lys96 and Asp64, a salt bridge between side chain of Lys60 and the free negatively charged carboxy C-terminus of αSyn(31-100) at Leu100 (Fig. 2B). This latter observation suggests that the larger interface of the tetrameric structure is enabled by the specific construct boundaries and may not be formed with full-length αSyn or other C-terminal truncations.

**Figure 2.**
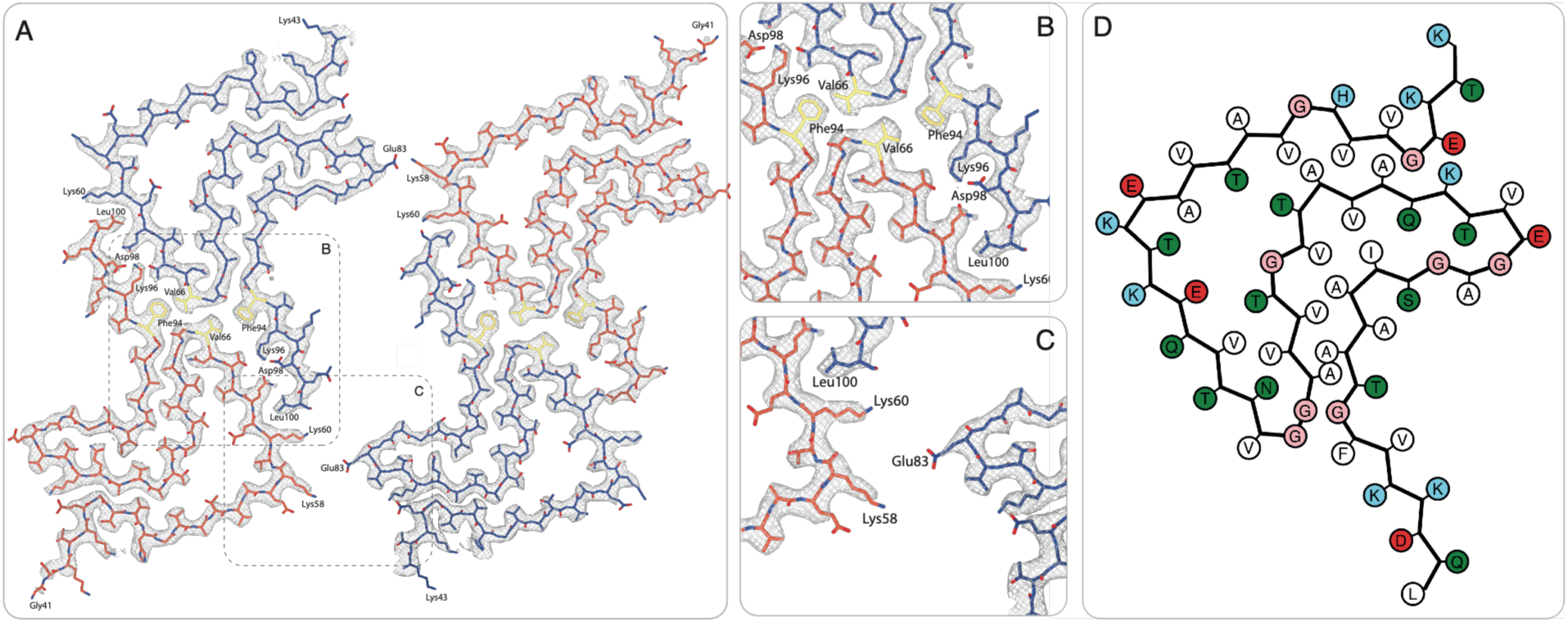
Atomic structure of type 1t polymorph of αSyn(31-100). A) Cryo-EM density map and atomic model of the tetrameric type 1 subtype formed at pH 6. Selected residues and intermolecular interface regions are indicated. B) Enlarged view of the large protofilament interface, C) enlarged view of the small protofilament interface. D) A schematic showing all amino acids involved in the fold.

At unphysiological pH 10 the new subtype of type 3F polymorph was identified for αSyn(31-100) amyloid fibrils (Fig. 3A-C). Representative cryo-EM data and validation are shown in the Supplementary (Fig. S4). The F interface of type 3F is relatively small and mediated by hydrophobic interactions between two Ala85 residues. A purely hydrophobic interface is atypical for αSyn fibrils, as protofilament interfaces are generally stabilized by one or two salt bridges. The absence of a salt bridge at the protofilament interface may be explained by the pKa of the lysine side chain (∼ 10.5), which results in partial loss of its positive charge under the aggregation conditions at pH 10. The observation of type 3F polymorph at high pH is unexpected, as type 3F polymorphs of full-length αSyn have previously been associated with acidic conditions (up to pH 6), above which type 2 polymorphs become dominant up to approximately pH 7 and then type 1 polymorphs at higher pH values.^15^ In this context, it is interesting to note that type 3 is observed for both protein constructs in the pH range close to their respective isoelectric points (pI) (i.e., pH ∼ 10 for αSyn(31-100), pI = 9.40, and pH ∼ 6 for full-length αSyn, pI = 4.67). This correlation may indicate that type 3F polymorph is preferred when the protein is in a net-neutral chemical configuration.

**Figure 3.**
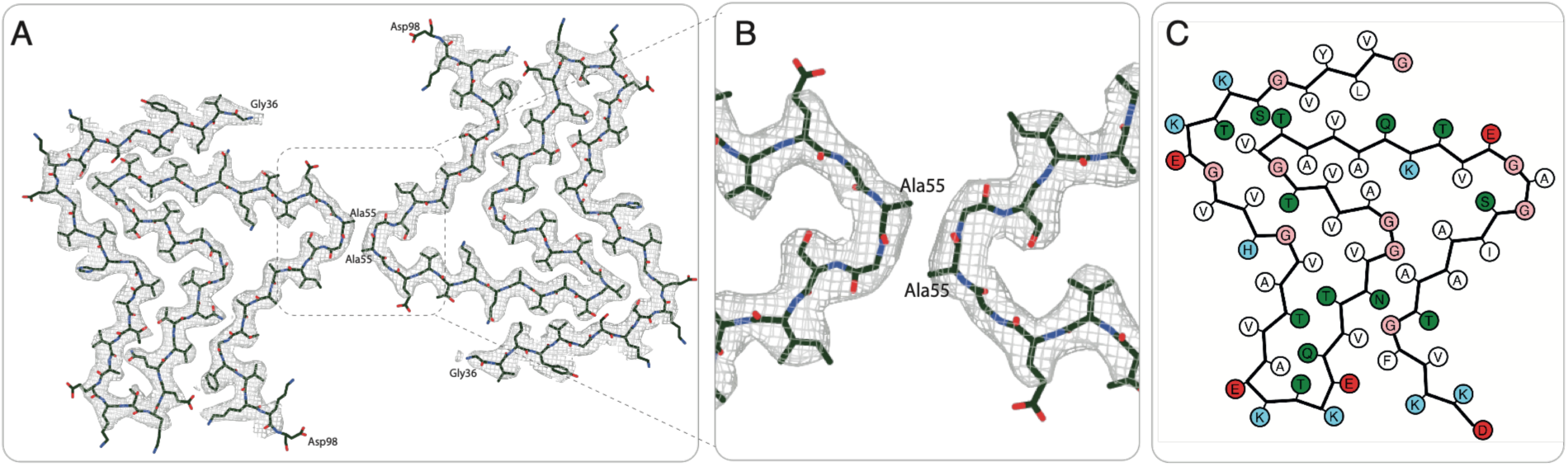
Atomic structure of type 3F polymorph of αSyn(31-100). A) Cryo-EM density map and atomic model of the type 3F formed at pH 10. Selected residues and intermolecular interface regions are indicated. B) Shows the enlarged view of the protofilament interface. C) A schematic showing all amino acids involved in the fold.

Within the intermediate pH range of approximately 6.5 - 9.5, the construct readily aggregated under multiple conditions; however, the resulting fibrils were not amenable to high-resolution cryo-EM structure determination.

Neutral and alkaline conditions yielded new subtypes of previously described type 1 and type 3 fibrils, whereas acidic conditions promoted formation of the novel type 10 polymorph.

Aggregation at pH 4 and 5 reproducibly promoted the formation of two previously unreported fibril folds, named type 10A and type 10B (Fig. 1, 4 and 5). Both structures were observed consistently across multiple aggregation conditions (Table 1). Whereas type 10A formed predominantly in the presence of divalent salts, type 10B was favored under monovalent salt conditions, suggesting that the ionic composition of the aggregation buffer influences polymorph selection.

**Figure 4.**
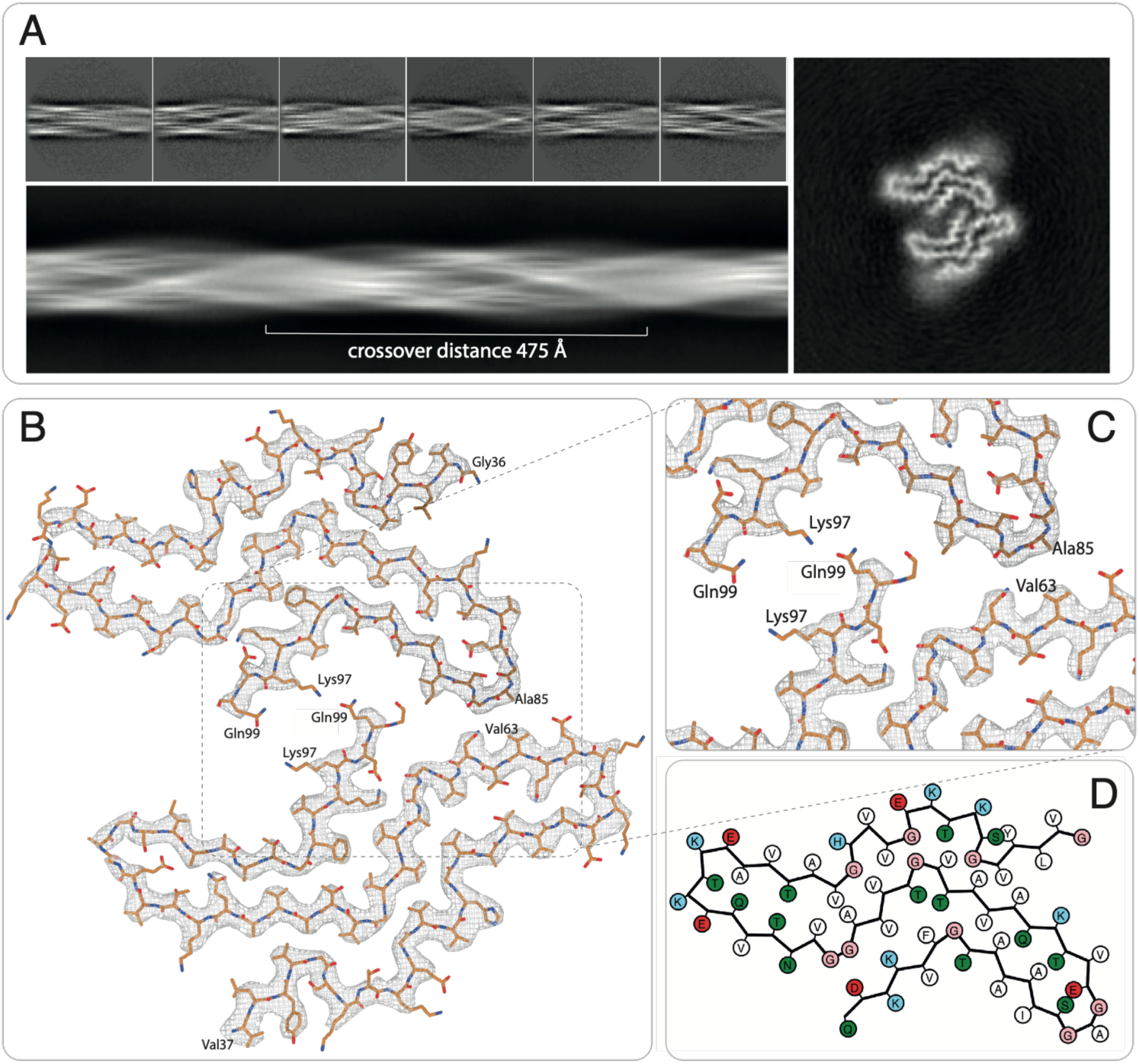
Structural characterization of the type 10A polymorph of αSyn(31-100). A) Representative cryo-EM 2D class averages of the fibril together with the projection of the reconstructed fibril volume. The fibril exhibits a crossover distance of 475 Å. The schematic on the right illustrates the protofilament arrangement and orientation of the N and C termini. B) Cryo-EM density map overlaid with the refined atomic model of the type 10A fibril cross section. C) Close-up of the interface in B with amino acid residues involved in intermolecular interactions indicated. D) A schematic showing all amino acids involved in the fold.

**Figure 5.**
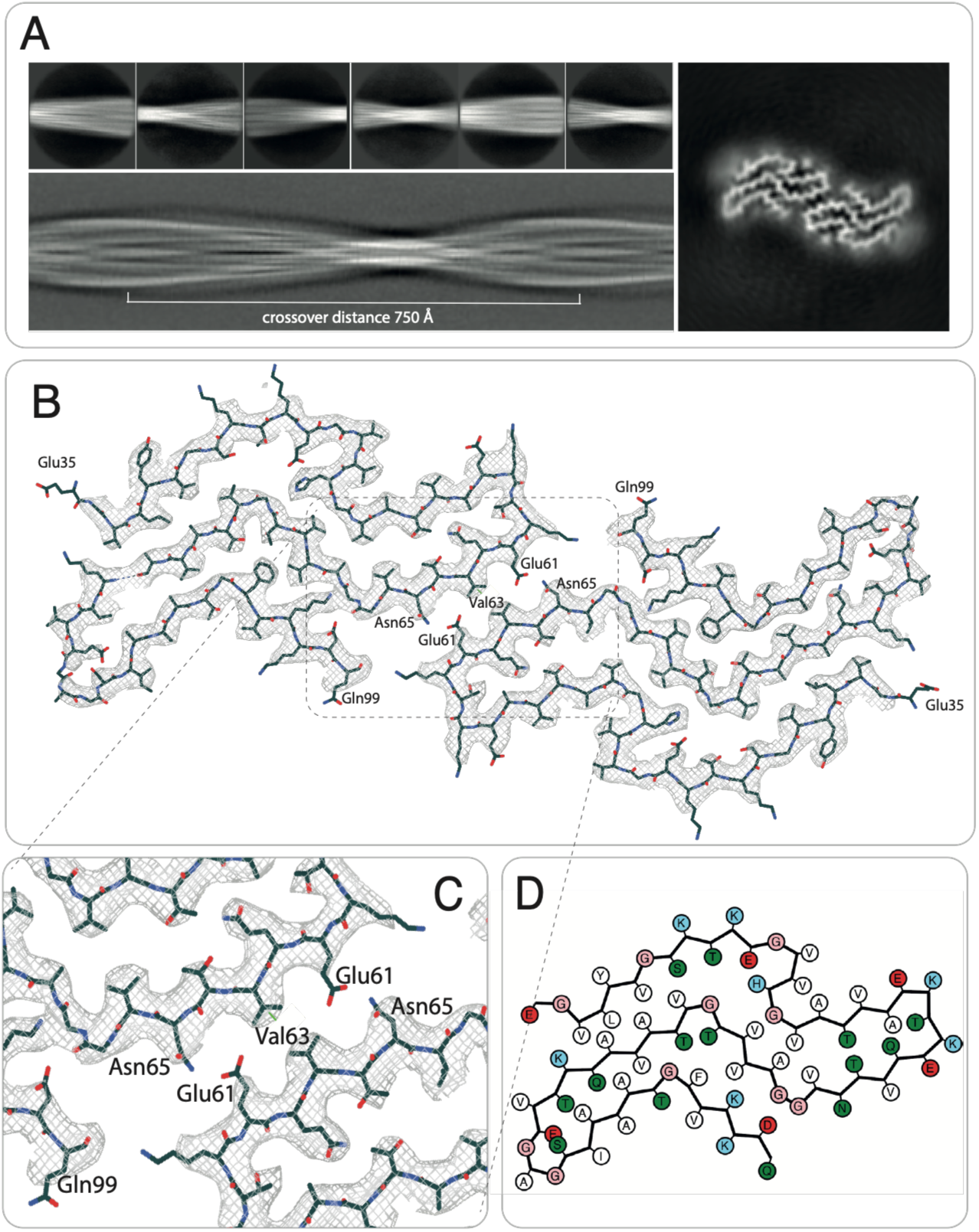
Structural characterization of the type 10B polymorph of αSyn(31-100). A) Representative 2D class averages, crossover distance, and a projection of the 3D fibril of the type 10B polymorph. B) Cryo-EM density map and atomic model of the type 10B polymorph. C) Enlarged view of the steric zipper interface in B, with interacting residues indicated. D) The schematics showing all amino acids involved in the fold.

### Structural characterization of the type 10 polymorph

Cryo-EM analysis revealed that type 10A (Fig. 4) and type 10B polymorph (Fig. 5) share a common protofilament fold but differ substantially in their protofilament interfaces. A structural overlay of the two protofilaments is shown in the Supplementary (Fig. S1). The two novel structures were reconstructed to final resolutions of 3.17 Å and 3.20 Å, respectively, according to the gold-standard FSC 0.143 criterion (Supplementary Figs. S5 and S6). While type 10A assembles into an asymmetric dimer, type 10B forms a symmetric dimeric fibril. The two polymorphs further differ in their helical symmetry and associated helical parameters. Type 10A exhibits a crossover distance of 475 Å, corresponding to a twist of -1.83° (Fig. 4A), whereas type 10B displays a much longer crossover distance of 750 Å, corresponding to a twist of -1.14° (Fig. 5A).

The novel type 10 protofilament has not been described previously and is characterized by several distinctive structural features. It is composed of eight β-strands with β1 comprising residues Val37-Val40 (having a β-bulge at position Val47/Leu48 in type 10B polymorph), β2 residues Lys43-Glu46, β3 residues Val52-Glu57, β4 residues Glu61-Val66, β5 residues Ala69-Thr72, β6 Val74-Glu83, β7 Ser87-Thr92, and β8 Phe94-Gln99 (Fig. 6A). With the exception of the C-terminal end of β3, all β-strands are flanked by Gly residues typically observed for amyloids (such as Gly73 and Gly84 at the borders of β6). The steric zipper is formed by a network of tightly packed strand-strand interactions comprising both hydrophobic and hydrophilic contacts. Hydrophobic interactions are observed between β1 and β6, involving Val37, Leu38, and Val40 on β1 and Val74, Val76, and Ala78 on β6. Additional partially hydrophobic contacts occur between β5 and β8 through Val70 and Phe94, and between β6 and β7 through Val77 and Ala89. In contrast, the interaction between β3 and β4 is predominantly hydrophilic and involves Thr54, Gln62, and Thr64. With respect to the influence of pH on the fibril structure, it is noteworthy that two glutamate residues, Glu46 and Glu83, are oriented toward the fibril interior. Glu46 is in proximity to His50, while Glu83 lies near Ser87. In both cases, no positively charged side chain is available for salt bridge formation, suggesting that these buried glutamate residues may be protonated, as has previously been shown for functional amyloids.^23^

**Figure 6.**
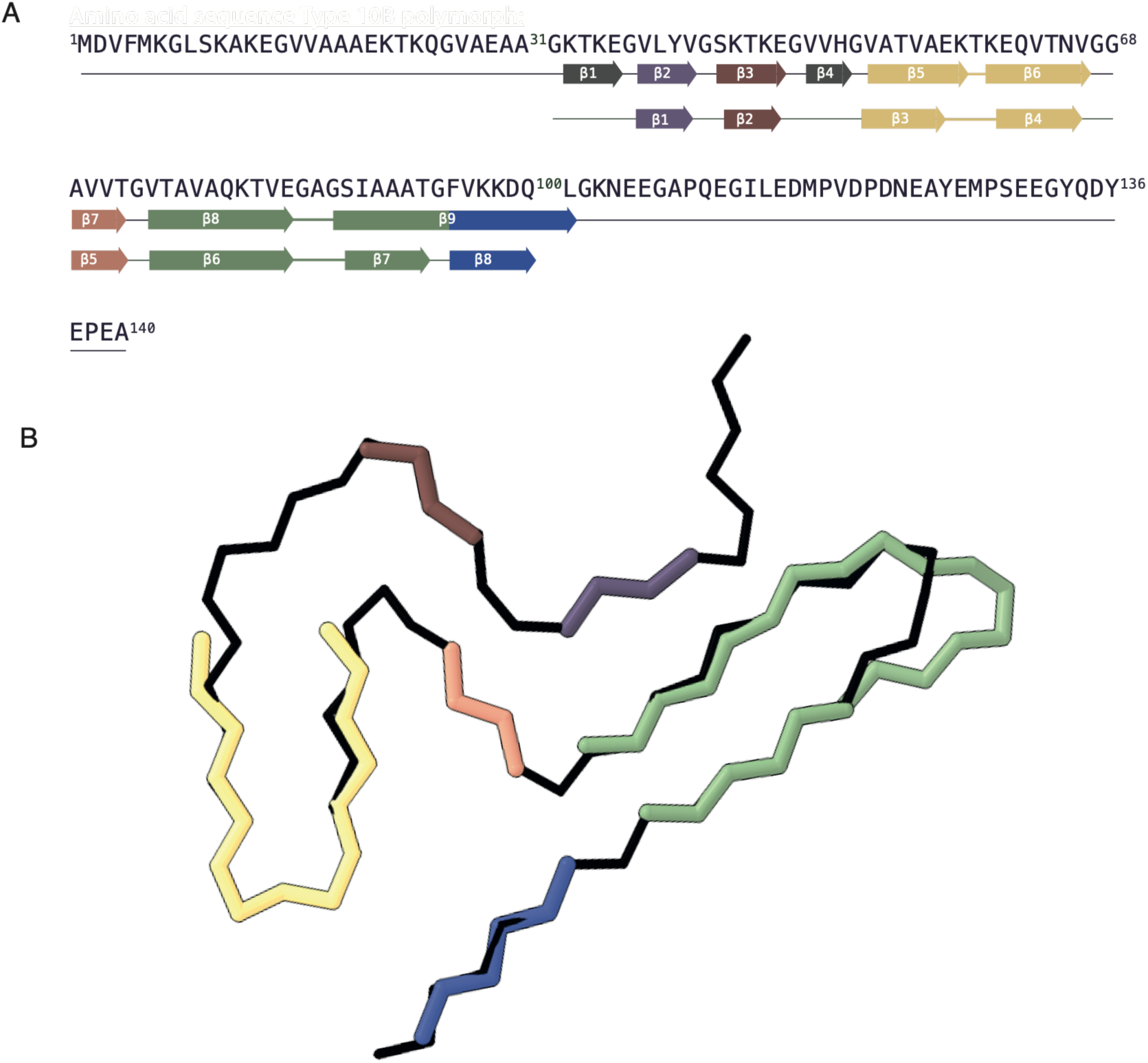
β-sheet structural comparison of the type 10 polymorph with the patient-derived PD-associated fold. A) Amino acid sequence of the full-length αSyn (top row) with the occurring β-sheets in the protofilament of the PD polymorph (middle row) and type 10 polymorph (bottom row). B) The β-sheets of the type 10 overlayed on the PD polymorph one by one to show the structural similarity.

This observation can be rationalized by the pKa of a free glutamate side chain (∼ 4.3), which is expected to be at least partially protonated under the acidic aggregation conditions (pH 4). Following this argumentation, it is not surprising that in contrast to the general finding of having several Lys-Glu salt bridges in αSyn polymorphs, they are missing in the type 10 polymorph. Only one Lys96-Asp98 salt bridge is present, attributed to the lower pKa of free Asp (∼ 3.9) compared to glutamate (∼ 4.3). A free Asp amino acid is negatively charged at pH 4 and thus the salt bridge formation can be rationalized.

The potential presence of protonated glutamate residues may also be relevant for the close packing of the type 10B polymorph interface, which is formed by Glu61, Val63, and Asn65 from one protofilament packing against the corresponding Glu61, Val63, and Asn65 residues of the opposing protofilament (Fig. 5C). In this arrangement, protonated Glu61 could participate in hydrogen-bonding interactions with Asn65, thereby contributing to stabilization of the interface. In striking contrast, the protofilament interface of type 10A polymorph is asymmetric and appears to be stabilized primarily by intermolecular hydrogen bonds between Lys97 and Gln99 residues. In addition, the asymmetric interface comprises Val63 from one protofilament and Ala85 from the opposing protofilament.

Additional unidentified densities were observed flanking the type 10B fibril but were absent in type 10A (Fig. 5A). Similar densities have been reported in multiple previously described αSyn fibrils and are often attributed to negatively charged ions.^16,24,25^ One possible explanation is the presence of phosphate species interacting with solvent exposed regions of the fibril surface. However, no obvious positively charged surface patch is located adjacent to these densities in the present structure. Furthermore, due to limited resolution in phosphate-free preparations adopting the same fold, the identity of these densities could not be resolved conclusively.

### Shared structural features of type 10 and the Parkinson’s Disease (PD) polymorph

The attempt to generate the PD polymorph using the αSyn(31-100) construct, that was successful in the case of Tau fibrils for the Alzheimer polymorph, failed for αSyn fibrils. The PD polymorph is composed of a single filament and has a distinct structure from type 10. However, a comparison between polymorph 10 with the patient-derived PD polymorph reveals structural similarities.

With the exception of the first two β-strands (β1 and β2), which participate in cofactor binding in the PD polymorph, the remaining β-strands (β3-β8) occupy similar positions in both the PD and type 10 structures (Fig. 6). Thus, at the secondary structure level the PD polymorph and polymorph 10 are similar. Moreover, the two characteristic loop regions 53 - 65 and 73 - 90 also resemble at the tertiary structural level (Fig. 6 Supplementary Fig. S2).

Together, these observations suggest that the combination of construct truncation and acidic aggregation conditions favors the formation of structural motifs associated with the PD polymorph.

Acidic conditions were motivated by physiological considerations: Intracellular compartments such as endosomes and lysosomes, which have pH values of around 4.5-5.5, and low pH gut regions have been proposed as potential sites of early αSyn aggregation. This raises the possibility that acidic environments may favor the formation of disease-relevant structural motifs.^26,27^

## Conclusion

The structural work on αSyn(31-100) amyloid fibrils demonstrates that truncation of the flexible terminal regions of αSyn influences fibril polymorphism across a broad range of physicochemical conditions. Systematic screening of pH and ionic environments revealed four previously unreported fibril structures, including novel subtypes of the established type 1 and type 3F polymorphs, as well as a new fibril fold that occurred reproducibly under acidic conditions (type 10). The structural analysis illustrates that the influence of truncation can arise from different sources. In the case of type 1 polymorphism, the direct influence of truncation on structure formation is evident in the formation of a salt bridge involving the C-terminal carboxylic acid at residue 100. In the case of the type 3F polymorph formed by αSyn(31-100) at pH 10, the influence of truncation may be rather at the overall protein charge level, under the assumption that type 3F polymorph is a preferred fold when the protein construct has a net zero charge, since its pI differs from the full-length αSyn.

Finally, the strong pH dependence of αSyn polymorph formation previously reported for full-length αSyn, which led to its characterization as a chameleon-like protein, is also recapitulated in the truncated αSyn(31-100) construct. Particularly noteworthy is the formation of the type 10 polymorph under acidic conditions (pH 4). Although type 10 differs substantially from the patient-derived PD polymorph, it reproduces several characteristic local structural motifs of the disease-associated polymorph. Notably, the formation of the type 10 polymorph at pH 4 was highly reproducible. One possible explanation for this is that, under acidic conditions, the protonation of glutamate side chains reduces the number of available Lys-Glu salt bridge interactions. This limits the range of accessible electrostatic interaction networks and constrains the structural landscape available during fibril assembly. This reduction in electrostatic complexity may favor the formation of the type 10 polymorph.

Furthermore, the emergence of type 10 under acidic conditions, together with its structural similarity to the PD polymorph derived from patients, further supports the hypothesis that acidic microenvironments may promote the formation of fibril structures relevant to the disease.

More broadly, our results demonstrate that rational construct design combined with systematic environmental screening and, finally, structure determination using cryo-EM represents a promising strategy for approaching disease-relevant αSyn conformations *in vitro*. It is of utmost importance to recapitulate the PD polymorph *in vitro*, since *in vitro* accessible disease-like fibrils will be valuable for mechanistic studies, structural comparisons, and the future development of conformation-specific diagnostic and therapeutic agents.

## Methods

### Recombinant protein expression and purification of αSyn

Recombinant N-terminally and C-terminally truncated, human αSyn was produced by expressing a pET-24a(+) plasmid in *Escherichia coli* BL21(DE3*). The construct was cloned into a pET-24a(+) vector purchased from GenScript with N-terminal His-Tag and a TEV site, cleaving between Q/G (G is the 31 amino acid).

In brief, a single colony from a fresh transformation was grown at 37 °C in 100 mL LB media containing 100 µg/mL kanamycin. The overnight primary growth culture was added to freshly prepared 1 L LB media containing 100 µg/mL kanamycin at a final concentration of 2.5% (vol/vol) and cultured at 37 °C with continuous agitation until the O.D. reached 0.8. Protein overexpression was initiated by adding IPTG at a final concentration of 1 mM. Thereafter, cells were grown at 37 °C for more than 4 h, harvested by centrifugation at 3000 g and lysed by three passes over a M110S microfluidizer with a Z-type interaction chamber at an input processing pressure of 45 psi. Purification of the lysate supernatant was performed using His tag affinity chromatography on a HisTrap FF column (Cytiva). The His tag was subsequently removed by overnight cleavage with TEV protease at a 200:1 protein to protease molar ratio. Following cleavage, the sample was applied stepwise (5 mL) onto a Ni NTA resin column to remove the cleaved His tag. The pure protein was dialyzed extensively against 10 mM ammonium acetate at pH 7, lyophilized, and stored at -20 °C until further use.

### Aggregation

Lyophilized αSyn was dissolved in PBS (pH above 6), 0.1 M citrate-phosphate or 0.1 M phosphate-borate (pH 10) buffer prepared from commercially available tablets (solution tampon phosphate tablet, pH 7.2-7.6 (1 tablet/200 mL), Sigma-Aldrich) and chemicals (Sodium citrate tribasic dihydrate ACS reagent grade, ≥99.0%, Sodium phosphate monobasic dihydrate purum p.a., crystallized, ≥99.0% (T), Sodium tetraborate decahydrate ReagentPlus, ≥99.5% by Sigma-Aldrich and di-Sodium hydrogen phosphate dihydrate ≥99.5% by VWR) and adjusted to the respective pH. Dissolved αSyn was separated from aggregates by passing it over two Vivaspin 500 (Sartorius) device (first with a MWCO filter of first 300 kDa and subsequently of 100 kDa) pre-rinsed with the same buffer and pH, yielding monomeric αSyn. Additives were added according to Table 1 and protein concentration was adjusted to 40 or 20 µM, respectively. 135 µl of the protein solution was incubated for 5 days in a 96 well UV-Star microplate (Greiner) and agitated for 1 min every 15 min at 37 °C in a BMG Pherastar plate reader at 250 rpm.

### Electron microscopy grid preparation and data collection

Cu R2/1 300 mesh grids (Quantifoils) were glow-discharged at -25 mA for 30 s, respectively. Freshly glow-discharged grids were used in a Vitrobot Mark IV (Thermo Fisher Scientific) with its chamber set at 100% humidity and at a temperature of 22°C. Fibrils (4 µl aliquots) were applied to the grid and blotted for 2.5-3.5 s after a 30-60 s wait time and subsequently plunge-frozen into a liquid ethane/propane mix. The grids were clipped and immediately used or stored in liquid nitrogen dewar.

Data acquisition was performed on a Titan Krios (Thermo Fisher Scientific) operating at 300 kV equipped with a Gatan Imaging Filter (GIF) with a 20 eV energy slit using Gatan’s K3 direct electron detector in counting mode. Movies were collected using EPU software (Thermo Fisher Scientific) at a magnification of ×130k and a dose rate of approximately 4-8 e/pixel/s and total dose ca. 50-75 e^-^/Å^2^.

### Image processing

Image processing and helical reconstruction were carried out with RELION 5.1, following the procedure for amyloid structures described by Scheres.^28^ RELION’s motion correction was used to correct for drift and dose-weighting, and CTF estimation was done using Ctffind4.1.^29^ Individual filaments were selected using Cryolo^30^, and segments were extracted in a 333 Å box with an inter-box distance of 33 or 66 Å and binned 4x to a pixel size of 2.6 Å for initial 2D classification (see Table S1 for details). To generate an initial model, *relion_helix_inidmod2d* was applied and the model could be used for the subsequent 3D refinement steps. For the final 3D refinements and CTF refinement, segments were extracted at a pixel size of 1.3 Å.

### Model building and refinement

Initial atomic models were generated from the postprocessed maps using ModelAngelo.^31^ The resulting models were manually inspected and corrected in COOT^32^, followed by refinement in ISOLDE.^33^ Refinement was performed on a 9-layer fibril model while applying symmetry restraints. The refined 9-layer model was additionally refinement in PHENIX^34^ to optimize atomic displacement parameters (B-factors). Finally, the two terminal layers at each end of the fibril, which frequently exhibit minor structural deviations, were removed and the central five layers were deposited in the PDB. Figures were prepared with Adobe Illustrator^35^ and UCSF Chimera^36^. Schematics showing all amino acid residues involved in the fold were produced using atom2svg.py.^37^

## Acknowledgements

We want to thank the staff at ScopeM ETHZ for their assistance with the Cryo-EM data collection analysis.

## Data availability

The reconstructed cryo-EM maps are deposited on the Electron Microscopy Data Bank with the accession codes EMD-58789 (type 1t), EMD-58790 (type 3F), EMD-58788 (type 10A), EMD-58787 (type 10B). The coordinates of the atomic model are deposited on the Protein Data Bank (PDB) under the accession code PDB 32BW (type 1t), 32BX (type 3F), 32BV (type 10A), 32BU (type 10B).

## Funding

We would like to thank the Synapsis foundation (2023-PI04) for financial support.

## Notes

### Competing Interest Statement

The authors have declared no competing interest.

## Bibliography

1. Lashuel, H. A. Rethinking protein aggregation and drug discovery in neurodegenerative diseases: Why we need to embrace complexity? Curr Opin Chem Biol 64, 67–75 (2021).

2. Goedert, M., Jakes, R. & Spillantini, M. G. The Synucleinopathies: Twenty Years On. Journal of Parkinson’s Disease 7, S51–S69 (2017).

3. Riek, R. The Three-Dimensional Structures of Amyloids. Cold Spring Harb Perspect Biol 9, a023572 (2017).

4. Ke, P. C. et al. Half a century of amyloids: past, present and future. Chem. Soc. Rev. 49, 5473–5509 (2020).

5. Xu, C. K. et al. α-Synuclein oligomers form by secondary nucleation. Nat Commun 15, 7083 (2024).

6. Riek, R. & Eisenberg, D. S. The activities of amyloids from a structural perspective. Nature 539, 227–235 (2016).

7. Goedert, M. Alpha-synuclein and neurodegenerative diseases. Nat Rev Neurosci 2, 492– 501 (2001).

8. Lashuel, H. A., Overk, C. R., Oueslati, A. & Masliah, E. The many faces of α-synuclein: from structure and toxicity to therapeutic target. Nat Rev Neurosci 14, 38–48 (2013).

9. Sawaya, M. R., Hughes, M. P., Rodriguez, J. A., Riek, R. & Eisenberg, D. S. The expanding amyloid family: Structure, stability, function, and pathogenesis. Cell 184, 4857– 4873 (2021).

10. Lövestam, S. et al. Seeded assembly *in vitro* does not replicate the structures of α-synuclein filaments from multiple system atrophy. FEBS Open Bio 11, 999–1013 (2021).

11. Yang, Y. et al. Structures of α-synuclein filaments from human brains with Lewy pathology. Nature 610, 791–795 (2022).

12. Schweighauser, M. et al. Structures of α-synuclein filaments from multiple system atrophy. Nature 585, 464–469 (2020).

13. Lashuel, H. A. Cracking the code of native amyloid fibrils: advances and next steps to enable pathology-informed therapeutic and diagnostic. Nat Struct Mol Biol 33, 567–576 (2026).

14. Stahlberg, H. & Riek, R. Structural strains of misfolded tau protein define different diseases. Nature 598, 264–265 (2021).

15. Frey, L. et al. On the pH-dependence of α-synuclein amyloid polymorphism and the role of secondary nucleation in seed-based amyloid propagation. eLife 12, (2024).

16. Guerrero-Ferreira, R. et al. Cryo-EM structure of alpha-synuclein fibrils. eLife 7, e36402 (2018).

17. Seuring, C. et al. Amyloid Fibril Polymorphism: Almost Identical on the Atomic Level, Mesoscopically Very Different. J. Phys. Chem. B 121, 1783–1792 (2017).

18. Frey, L. et al. On the Polymorph-Selection Determinants of α-Synuclein Amyloid Fibrils Studied at Atomic Resolution. 2026.07.13.737774 Preprint at 10.64898/2026.07.13.737774 (2026).

19. McGlinchey, R. P., Ramos, S., Dimitriadis, E. K., Wilson, C. B. & Lee, J. C. Defining essential charged residues in fibril formation of a lysosomal derived N-terminal α-synuclein truncation. Nat Commun 16, 3825 (2025).

20. Sorrentino, Z. A. & Giasson, B. I. The emerging role of α-synuclein truncation in aggregation and disease. Journal of Biological Chemistry 295, 10224–10244 (2020).

21. McGlinchey, R. P., Ni, X., Shadish, J. A., Jiang, J. & Lee, J. C. The N terminus of α-synuclein dictates fibril formation. Proceedings of the National Academy of Sciences 118, e2023487118 (2021).

22. Lövestam, S. et al. Assembly of recombinant tau into filaments identical to those of Alzheimer’s disease and chronic traumatic encephalopathy. eLife 11, e76494 (2022).

23. Nespovitaya, N. et al. Dynamic Assembly and Disassembly of Functional β-Endorphin Amyloid Fibrils. J. Am. Chem. Soc. 138, 846–856 (2016).

24. Guerrero-Ferreira, R. et al. Two new polymorphic structures of human full-length alpha-synuclein fibrils solved by cryo-electron microscopy. eLife 8, e48907 (2019).

25. Lövestam, S. et al. Seeded assembly in vitro does not replicate the structures of α-synuclein filaments from multiple system atrophy. FEBS Open Bio 11, 999–1013 (2021).

26. Stephens, A. D., Zacharopoulou, M. & Kaminski Schierle, G. S. The Cellular Environment Affects Monomeric α-Synuclein Structure. Trends in Biochemical Sciences 44, 453–466 (2019).

27. Park, H., Kam, T.-I., Dawson, V. L. & Dawson, T. M. α-Synuclein pathology as a target in neurodegenerative diseases. Nat Rev Neurol 21, 32–47 (2025).

28. Scheres, S. H. W. Amyloid structure determination in RELION-3.1. Acta Cryst D 76, 94– 101 (2020).

29. Rohou, A. & Grigorieff, N. CTFFIND4: Fast and accurate defocus estimation from electron micrographs. Journal of Structural Biology 192, 216–221 (2015).

30. Wagner, T. et al. SPHIRE-crYOLO is a fast and accurate fully automated particle picker for cryo-EM. Commun Biol 2, 218 (2019).

31. Jamali, K. et al. Automated model building and protein identification in cryo-EM maps. Nature 628, 450–457 (2024).

32. Emsley, P. & Cowtan, K. Coot: model-building tools for molecular graphics. Acta Cryst D 60, 2126–2132 (2004).

33. Croll, T. I. ISOLDE: a physically realistic environment for model building into low-resolution electron-density maps. Acta Cryst D 74, 519–530 (2018).

34. Liebschner, D. et al. Macromolecular structure determination using X-rays, neutrons and electrons: recent developments in Phenix. Acta Cryst D 75, 861–877 (2019).

35. McNicholas, S., Potterton, E., Wilson, K. S. & Noble, M. E. M. Presenting your structures: the *CCP* 4 *mg* molecular-graphics software. Acta Crystallogr D Biol Crystallogr 67, 386– 394 (2011).

36. Pettersen, E. F. et al. UCSF Chimera—A visualization system for exploratory research and analysis. Journal of Computational Chemistry 25, 1605–1612 (2004).

37. Nakane, T. atom2svg. https://doi.org/10.5281/zenodo.4090925 (2020) doi:10.5281/zenodo.4090925.

