## Supplementary Data for "Cryo-EM structures of α-Synuclein(31-100) amyloid fibrils reveal disease-like structural motifs without reproducing the Parkinson’s Disease polymorph"

**Table 2: Cryo-EM structure determination information**

| Polymorph | 10A | 10B | 1t | 3F |
|---|---|---|---|---|
| **Data Collection** | | | | |
| Pixel size [Å] | 0.65 | 0.65 | 0.65 | 0.65 |
| Defocus range [μM] | -0.8 to -2.2 | -0.8 to -2.2 | -0.8 to -2.2 | -0.8 to -2.2 |
| Voltage [kV] | 300 | 300 | 300 | 300 |
| Number of frames | 40 | 40 | 40 | 40 |
| Total dose [$e^-$/ $Å^2$] | 67.2 | 56.7 | 54.8 | 56.7 |
| **Reconstruction** | | | | |
| Box width [pixels] | 256 | 256 | 256 | 256 |
| Inter-box distance [Å] | 66 | 66 | 33 | 66 |
| Reconstructed Pixel size [Å] | 1.3 | 1.3 | 1.3 | 1.3 |
| Number of Micrographs | 690 | 2153 | 1’534 | 355 |
| Extracted segments | 179’411 | 778’858 | 551’540 | 86’412 |
| Extracted segments after 2D classification | 34’011 | 56’355 | 36’809 | 27’507 |
| 3D refinement resolution [Å] | 3.54 | 3.66 | 3.03 | 3.29 |
| Final resolution [Å] | 3.17 | 3.20 | 2.68 | 2.92 |
| Estimated map sharpening B-factor [Å] | - 61.21 | - 100.25 | - 46.21 | - 51.97 |
| Axial symmetry | C1 | C1 | C1 | Pseudo $C2_1$* |
| Helical rise [Å] | 4.72 | 4.78 | 4.75 | 2.39 |
| Helical twist [º] | -1.83 | -1.14 | -1.21 | 179.34 |

*Pseudo $C2_1$ symmetry: a two-start C1 helix wrapped around each other with a relative shift in helical rise of about half of 4.75 Å.

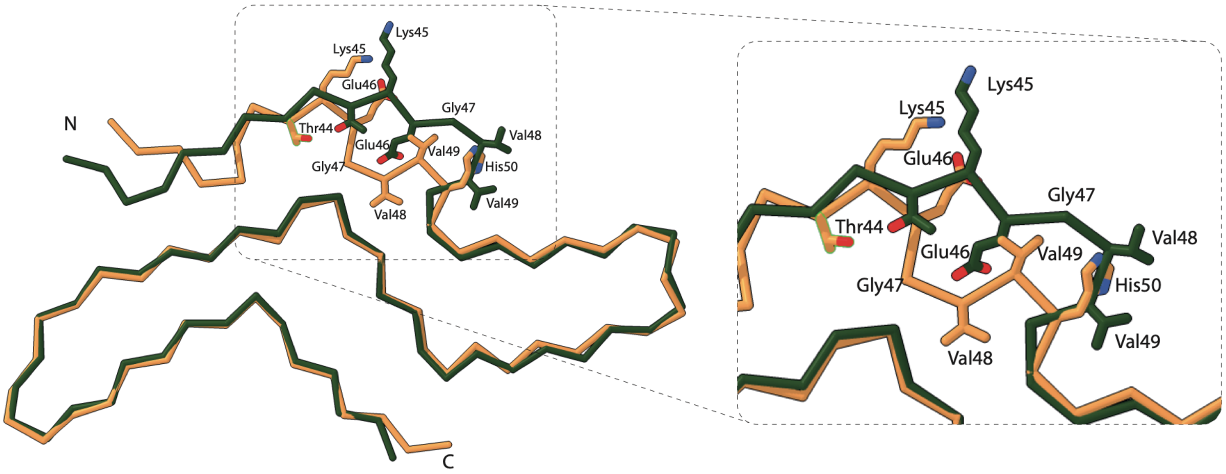


**Figure S1. Structural comparison of the type 10A polymorph and 10B.** Structural overlays of the type 10A and 10B protofilament. The overall protofilament fold is identical, with structural differences confined to residues 44 – 50, where distinct side-chain orientations and backbone conformations are observed.

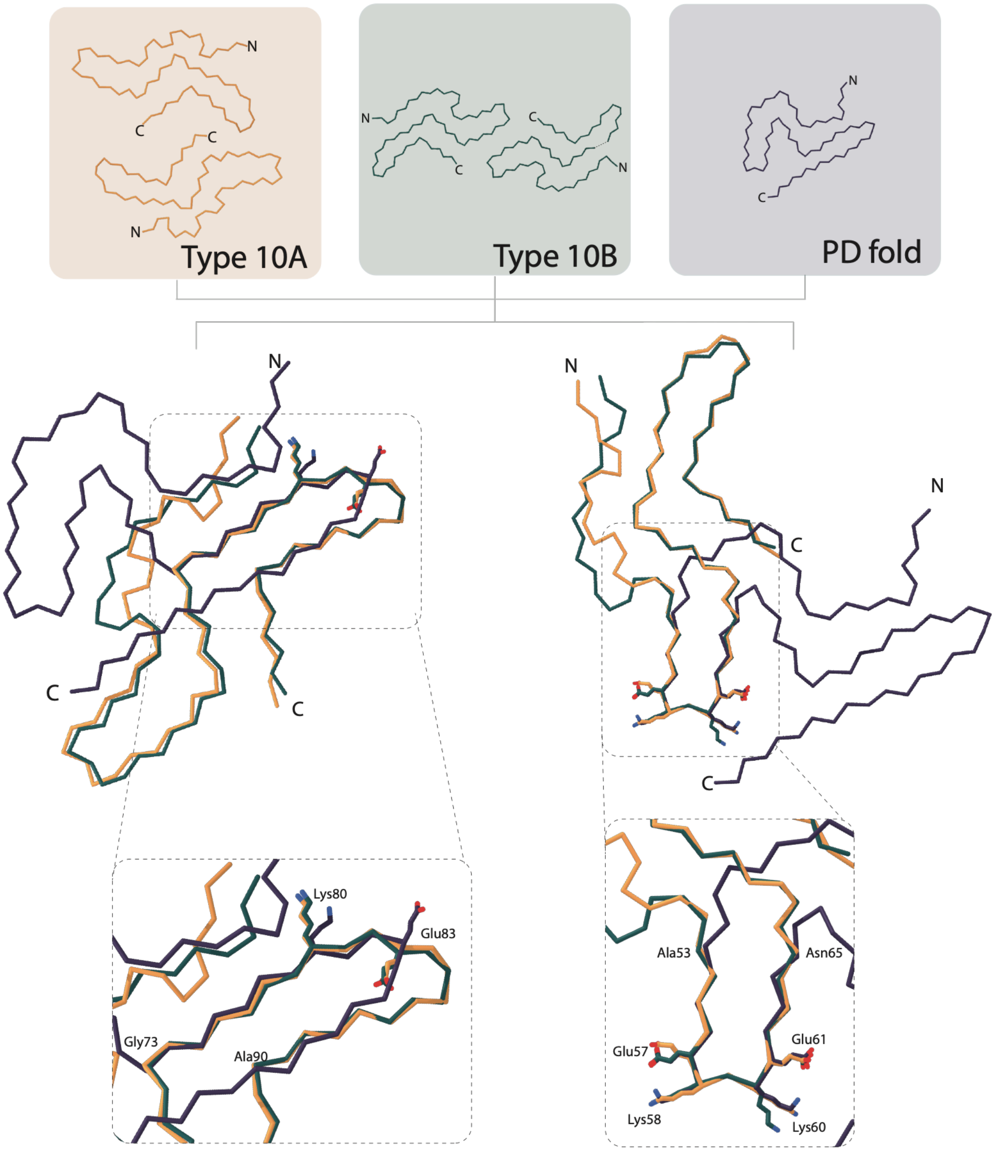


**Figure S2. Structural comparison of the type 10 polymorph with the patient-derived PD polymorph.** A) Schematic representation of the type 10A, type 10B and PD polymorph. B) Structural overlays of the type 10 polymorphs with the PD polymorph. Conserved backbone conformations are observed particularly within residues 53 - 65 and 73 - 90, where the type 10 structures adopt loop architectures highly similar to those of the patient-derived polymorph.

A

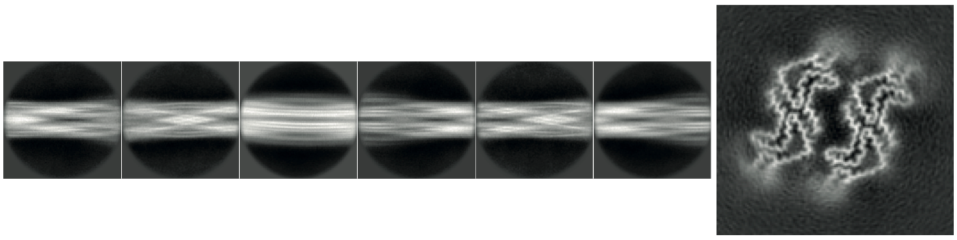

B

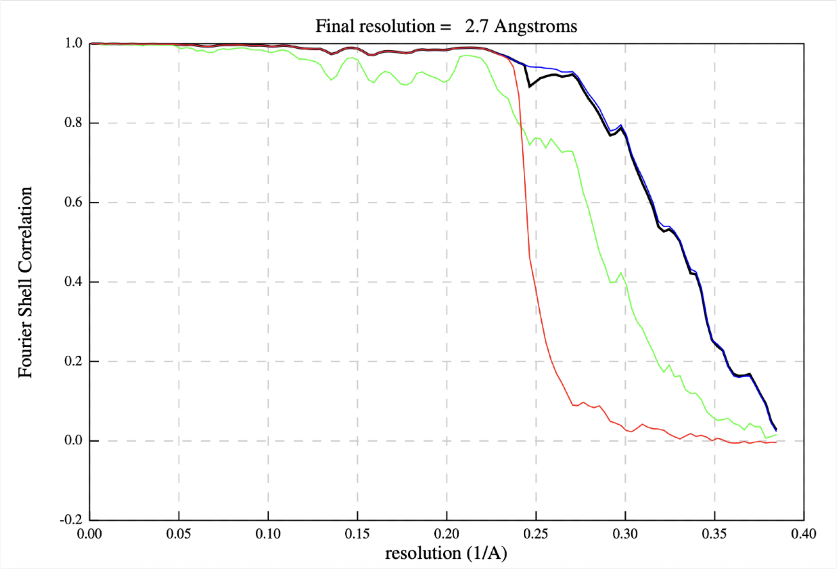


C

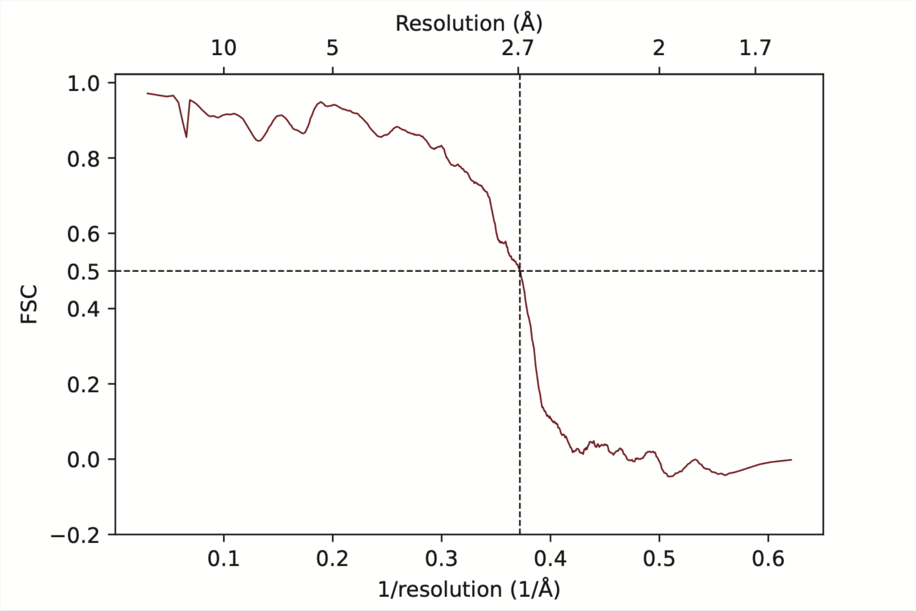


**Figure S3. 2D classes, slice through the 3D map and FSC curves for type 1t polymorph.** A) Six representative 2D classes used for the 3D reconstruction. On the right, a slice through the 3D reconstructed map is shown. B) FSC curve of the postprocessed map from RELION. Red curve: phase randomized masked map, green: unmasked map, blue: masked map. C) FSC curve of the model-map from PHENIX.

A

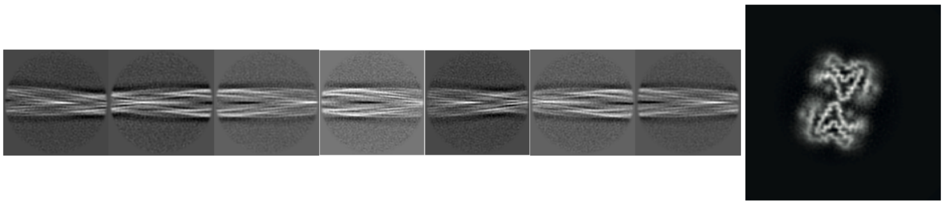

B

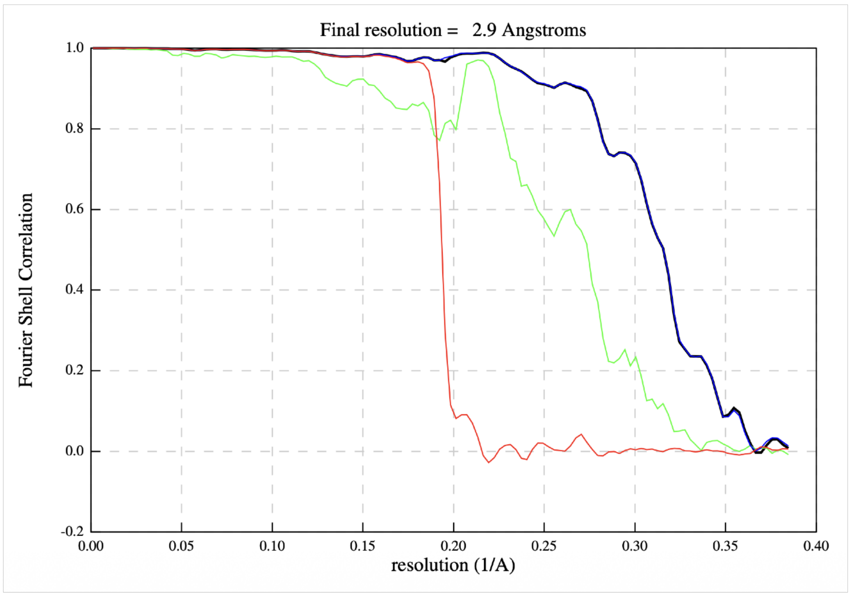


C

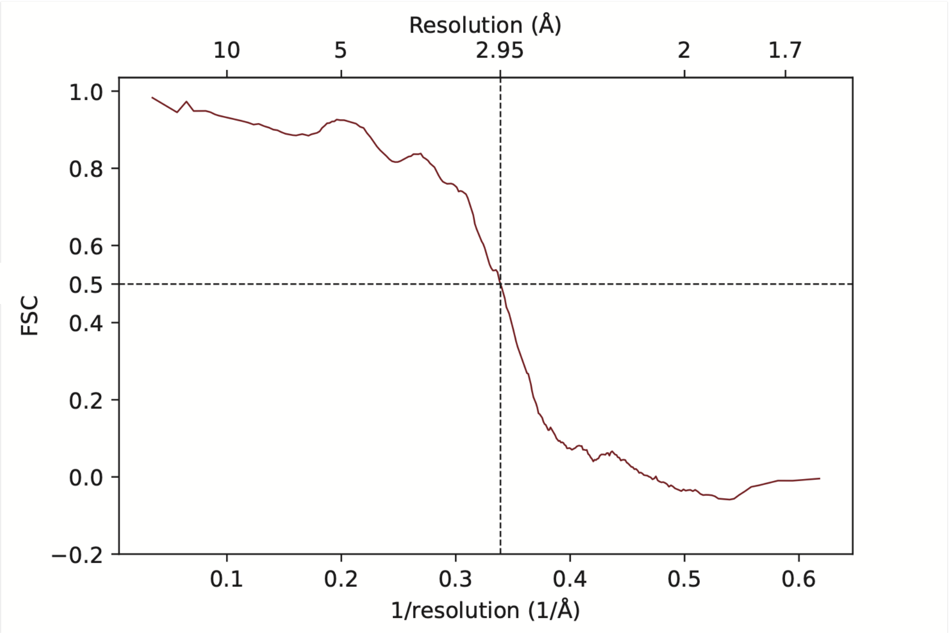


**Figure S4. 2D classes, slice through the 3D map and FSC curves for type 3F polymorph.** A) Seven representative 2D classes used for the 3D reconstruction. On the right, a slice through the 3D reconstructed map is shown. B) FSC curve of the postprocessed map from RELION. Red curve: phase randomized masked map, green: unmasked map, blue: masked map. C) FSC curve of the model-map from PHENIX.

A

B

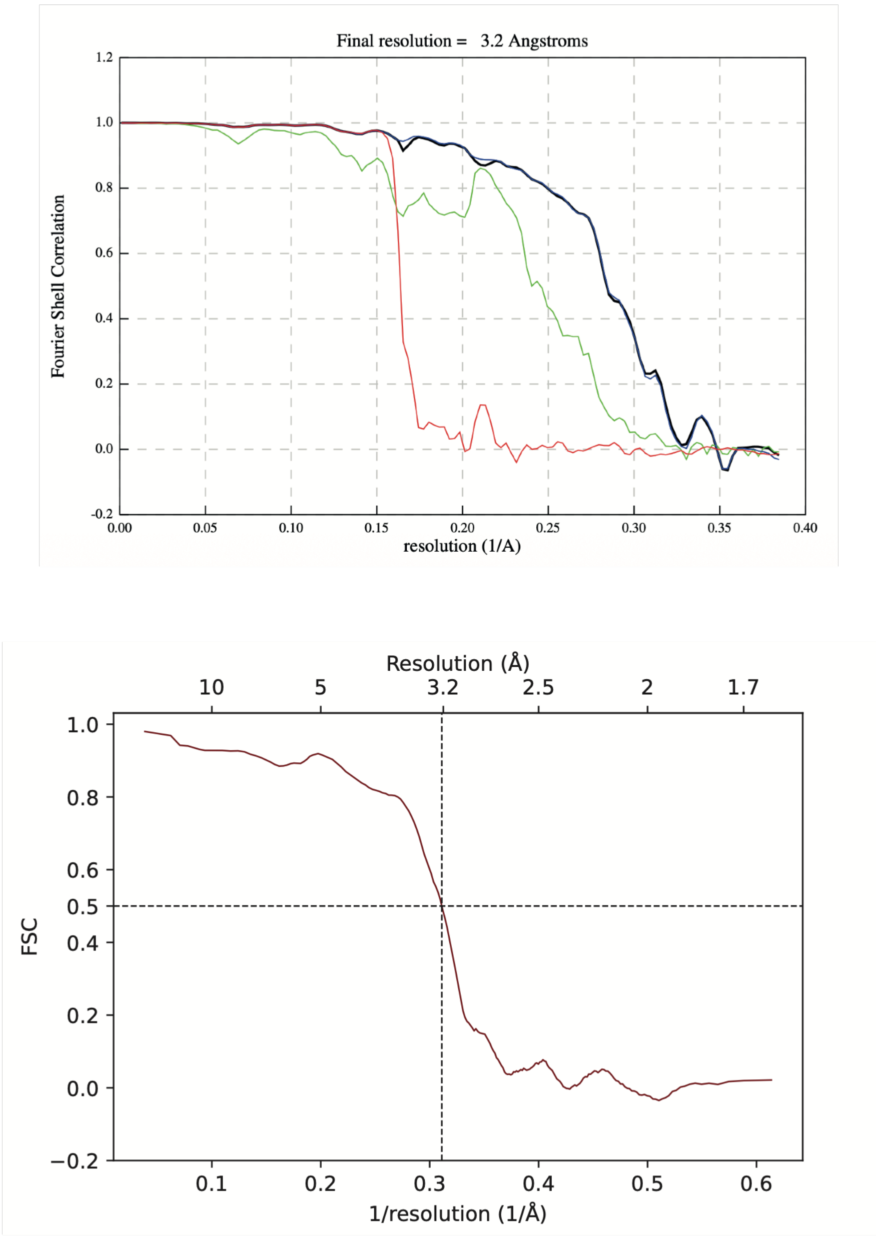


**Figure S5. FSC curves for type 10A polymorph.** A) FSC curve of the postprocessed map from RELION. Red curve: phase randomized masked map, green: unmasked map, blue: masked map. B) FSC curve of the model-map from PHENIX.

A

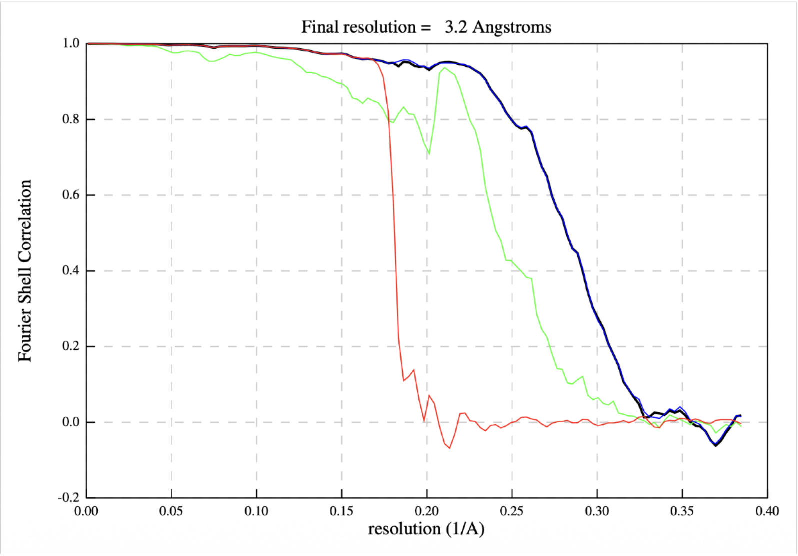


B

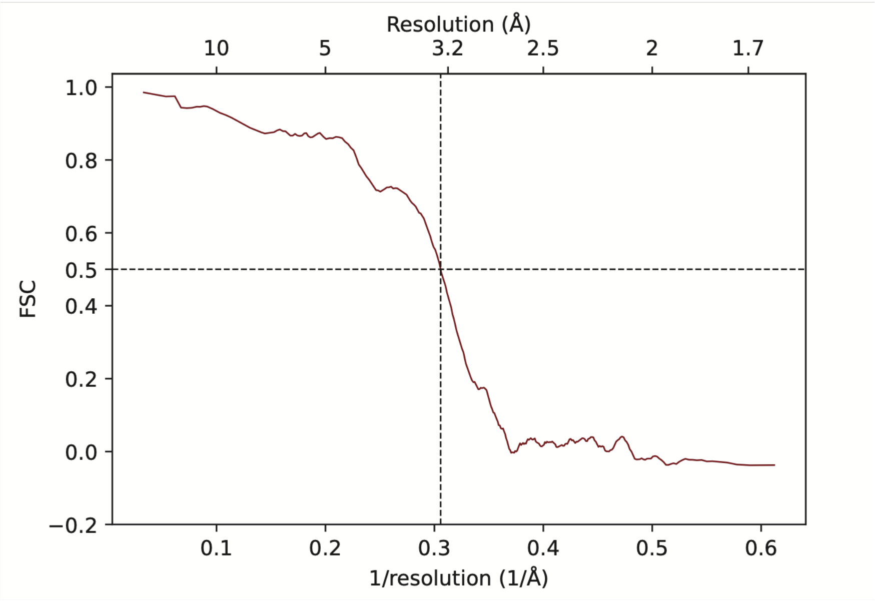


**Figure S6**. **FSC curves for type 10B polymorph.** A) FSC curve of the postprocessed map from RELION. Red curve: phase randomized masked map, green: unmasked map, blue: masked map. B) FSC curve of the model-map from PHENIX.
